# Netrin-1 inhibition does not attenuate cancer-induced bone pain in three translational models

**DOI:** 10.64898/2026.07.10.733710

**Authors:** Chelsea Hopkins, Mie Brandt Lassen, Louise Ploug Hansen, Yingyu Tang, Edward Ciputra, Theis Lund Jørgensen, Mads Haaber Christensen, Cecilie Louise Pedersen, Camilla I Svensson, Ming Ding, Rasmus S. Pedersen, Nicholas Willumsen, Anne-Marie Heegaard

## Abstract

Cancer-induced bone pain (CIBP) occurs in a majority of patients when primary or metastatic cancer develops within the bone. This pain has a significant impact on quality of life, yet there are limited effective treatment options available. Nerve sprouting is a complex mechanism that has been implicated in CIBP. Netrin-1 is a neuronal guidance molecule that is produced by numerous cell types, including cancer cells. In this study we aimed to determine whether netrin-1 inhibition (with NP137 – a humanized IGg1 monoclonal antibody) could ameliorate nerve sprouting, and nociception by extension, in three models of CIBP – osteosarcoma, metastatic breast cancer, and metastatic prostate cancer. Sustained administration of NP137 failed to produce an anti-nociceptive effect in these models, but a delayed onset was observed in the osteosarcoma model. NP137 did not produce a disease-modifying effect, as micro-computed tomography did not reveal reduced bone destruction in the NP137-treated groups. Additionally, there was no nerve fibre density reduction in any of the groups at the late-stage of the disease, suggesting that nerve sprouting occurs in early- to mid-stage CIBP development. Investigation of NP137 exposure indicated that serum levels of NP137 were comparable between the sham and cancer groups. Our study indicates that netrin-1 may play a role in early-stage CIBP development, but inhibition of this mechanism does not produce robust anti-nociception.

## 3. Introduction

Cancer-induced bone pain (CIBP) occurs when primary bone cancer, or metastatic cancer develops within the bone [1]. Among patients with bone involvement, breakthrough CIBP is common, and approximately half of patients experience moderate to severe pain [2]. Patients with CIBP experience significant physical impairment and their overall quality of life is significantly impaired and correlated with pain severity [3]. Yet, treatment options for CIBP rely on the World Health Organization principles set out in the 1986 Analgesic Ladder (updated in 2023), in which opioids remain a mainstay treatment for moderate to severe pain [4]. Improvements in cancer survival have led to an increasing number of patients living with CIBP for extended periods. Consequently, long-term opioid therapy is often required, despite its association with tolerance and severe adverse effects including treatment-limiting constipation [5], [6].

There are numerous mechanisms that may contribute to CIBP, including inflammation, nerve damage, and neurotrophic factors. These nociceptive mechanisms are mediated by the cancer cells, native bone tissue, and immune cells. Nerve sprouting is a complex phenomenon that differs between bone regions [7]. Nerve growth factor (NGF) is a well-studied protein produced by numerous cell types, promoting nerve growth and neuronal sensitivity. Previous CIBP studies have indicated that tropomyosin receptor kinase A (NGF receptor) inhibition in mouse models of osteosarcoma and prostate cancer leads to reduced nociception and reduces nerve presence in the periosteum [8], [9] and in a preclinical model of Lewis Lung carcinoma induced bone pain, NGF inhibition with a humanized monoclonal antibody showed an anti-nociceptive effect [10]. However, the clinical development of anti-NGF therapeutic antibodies has been limited by safety concerns, particularly the occurrence of rapidly progressive osteoarthritis in some patients, preventing regulatory approval [11]. Nevertheless, NGF is not the only molecular mechanism that controls nerve sprouting.

Netrin-1 is a neuronal guidance molecule, coordinating nerve growth direction by binding to deleted in colorectal carcinoma (DCC) or uncoordinated 5 (UNC5) receptors [12], but more recent studies have shown that netrin-1 may be implicated in painful conditions and cancer progression. Netrin-1 is suggested to contribute to nociception in neuropathic pain models [13], [14], [15], fibromyalgia [16], discogenic pain [17], endometriosis [18], central-mediated mechanical hypersensitivity [19], osteoarthritis [20], rheumatoid arthritis [21], and CIBP [22], [23]. Zhu et al. demonstrated that osteoclasts produced netrin-1, which contributed to *in vitro* sprouting and nociception in a mouse model of osteoarthritis [20] and Rudjito et al. demonstrated that netrin-1 inhibition was effective in reducing nociception in a mouse model of late-stage rheumatoid arthritis [21]. Similarly, Gong et al. demonstrated that DCC inhibition reduced nociception in a rat model of CIBP [22], and Diaz-delCastillo et al. demonstrated that netrin-1 inhibition produced a transient anti-nociceptive effect in early-stage CIBP [23]. Rudjito et al. and Diaz-delCastillo et al. both used a netrin-1 inhibitor, NP137, to produce anti-nociception in their respective models. NP137 is a monoclonal antibody that has also been shown to suppress cancer progression [24]. As such, netrin-1 blockade has the potential to attenuate nociception and prevent tumour progression within the bone.

Together, these studies suggest that NP137 treatment to attenuate netrin-1 may produce an anti-nociceptive effect in CIBP. This study aims to determine the analgesic efficacy of NP137 in three murine CIBP models – osteosarcoma, metastatic-like breast cancer, and metastatic-like prostate cancer.

## 4. Methods

### 4.1 Animals

All experiments were approved and conducted in accordance with the Danish Animal Experiments Inspectorate (Copenhagen, Denmark, 2020_15_0201_00439). Three mouse strains were used, each paired with its corresponding syngeneic cell line to ensure all models were immunologically compatible. Balb/cOlaHsd (female; n = 48) was used for the breast cancer model, C57BL/6JOlaHsd (male; n = 48) was used for the prostate cancer model, and C3H/HeNHsd (female; n = 50) was used for the osteosarcoma model. Mice were group-housed in a specific pathogen-free facility at the University of Copenhagen. They had 12-hour light/dark cycles, ad libitum access to water and standard chow, and provided with bedding, nesting material, and enrichment items. Mice were excluded from nociceptive behaviour analysis if they did not develop tumours after being injected with cancer cells, confirmed through x-ray imaging.

### 4.2 Tumour cells

NCTC 2472 osteosarcoma cells (CCL-11, American Type Culture Collection, Manassas, VA, USA) were cultured in NCTC 135 medium (Gibco, Carlsbad, CA, USA) with 10% horse serum (Gibco, Carlsbad, CA, USA, Lot: 2593036). RM-1 prostate cancer cells (CRL-3310, American Type Culture Collection, Manassas, VA, USA) were culture in Dulbecco’s Modified Eagle Medium (Gibco, Carlsbad, CA, USA) with 10% fetal bovine serum (Thermo Fisher Scientific, lot: 2556826RP, MA, USA), and 1% penicillin-streptomycin (Gibco, Carlsbad, CA, USA). 4T1-Luc2 mammary carcinoma cells (CRL-2539-LUC2, American Type Culture Collection, Manassas, VA, USA) were cultured in RPMI 1640 medium (without phenol red, Gibco, Carlsbad, CA, USA), with 10% fetal bovine serum (Thermo Fisher Scientific, lot: 2556826RP; MA, USA), and 1% penicillin-streptomycin-glutamine (Gibco, Carlsbad, CA, USA). Cells were cultured for two weeks prior to cancer cell inoculation surgery. Cells were passaged when they reached 80% confluency, two days prior to surgery, and on the morning of the surgery. All cells are adherent and cultured in 75cm^2^ flasks (Corning, New York, USA) and treated with 0.05% trypsin-EDTA for four minutes at 37°C until cells released from the flask, after which, trypsin was quenched with complete medium. For surgery, cells were resuspended in 0.1M PBS at 10^7^ cells/ml (NCTC 2472 cells) or 1X HBSS (Gibco, Carlsbad, CA, USA) at 10^6^ cells/ml (RM-1 and 4T1-Luc2 cells).

### 4.3 Drugs

A ketamine/xylazine cocktail in sterile saline was freshly prepared prior to surgery (43 mg/kg Ketaminol, MSD Animal Health, Boxmeer, The Netherlands; 6 mg/kg Xylazine, Rompun Vet, Bayer, Germany). The cocktail was administered via intraperitoneal injection (i.p.) approximately 10-15 min prior to surgery. A cocktail of 85.5mg/kg ketamine and 12mg/kg xylazine in saline was freshly prepared for perfusions. During surgery, mice were maintained on 1-1.2% isoflurane (Attane Vet, ScanVet, Fredensborg, Denmark) until surgery was completed. Mice were placed on 2-2.5% isoflurane when surgical clips were removed (Day 1-4) and during x-ray imaging. Mice received carprofen (5 mg/kg Carprosan Vet, Dechra, Leverkusen, North Rhine-Westphalia, The Netherlands) immediately prior to surgery and on Day 1 via subcutaneous injection as a surgical analgesia.

NP137 (Netris Pharma, Lyon, France) is a monoclonal antibody against netrin-1 [24]. NP137 was administered i.p. with a loading dose of 20mg/kg on Day 4 after surgery and administered three times per week (from Day 5) thereafter at 10mg/kg (Figure 1A). The dose and schedule were in accordance with the current treatment approach at NETRIS Pharma and previous studies [23], [24]. Sterile saline was provided at a similar volume (5ul/g body weight) in the vehicle treated group. Etanercept (Erelzi, Sandoz, Basel, Switzerland) was administered at 30mg/kg i.p. once per week from Day 8. Experimenters were masked to the treatment groups until all analysis was completed.

**Figure 1:**
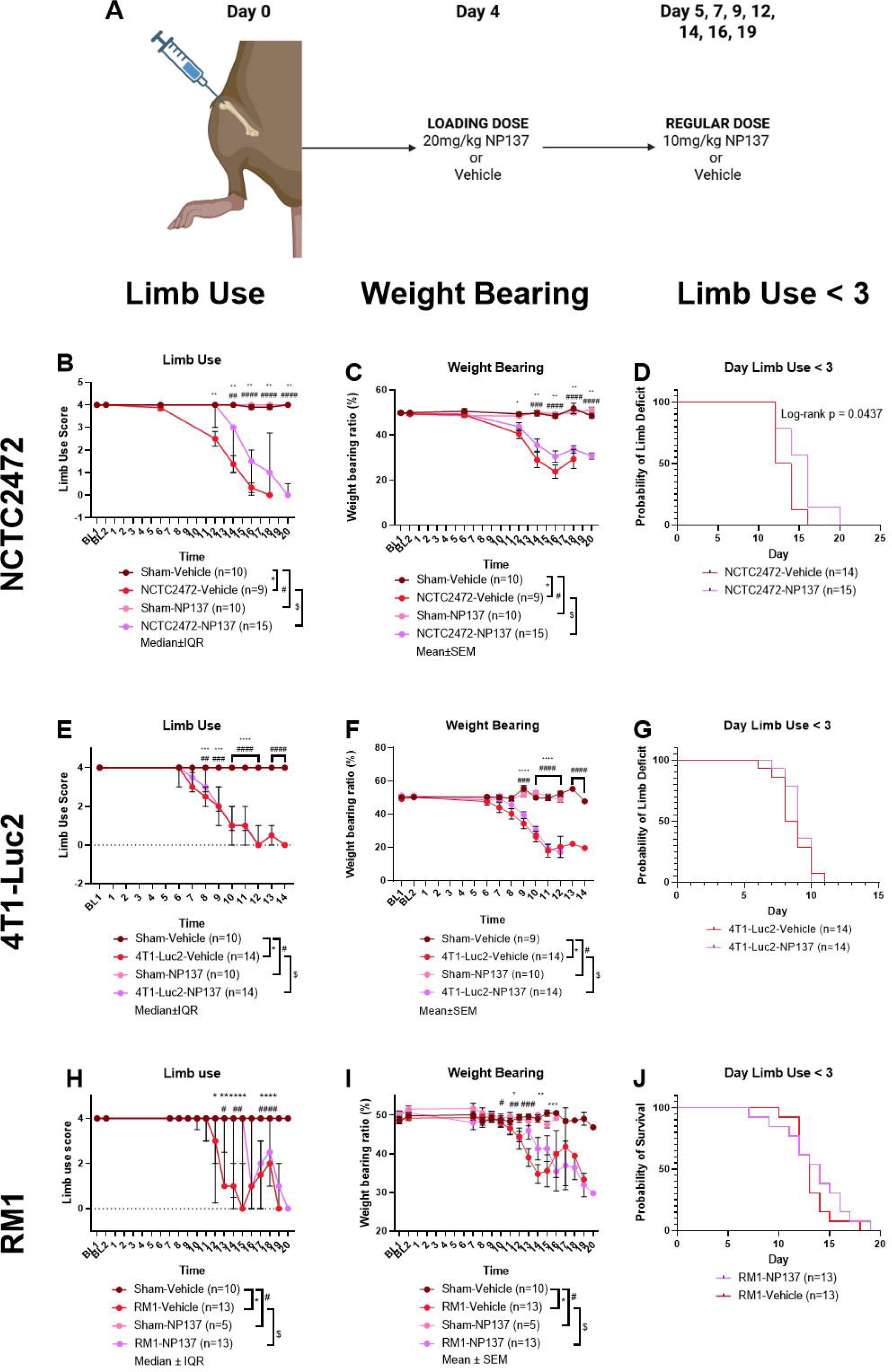
Behavioural analysis performed on three cancer-induced bone pain models using limb use and static weight bearing assessments. (A) Dosing diagram demonstrating cancer cell inoculation on day 0, a loading dose on day 4, and repeated doses thereafter; (B, C, D) behavioural outcomes for NCTC2472 model; (E, F, G): behavioural outcomes for 4T1-Luc2 model; (H, I, J): behavioural outcomes for RM1 model; (B, E, H): limb use assessment outcomes, with a two-way ANOVA assessment with Tukey Test and presented as median±IQR; (C, F, I) weight bearing assessment outcomes, with a two-way ANOVA assessment with Tukey Test and presented as mean±SEM; (D, G, J): survival plot displaying day at which a limb use score below 3 was achieved (log-rank test). (BL: baseline; n: number; IQR: interquartile range; SEM: standard error of the mean; ANOVA; analysis of variance; *: p < 0.05; **: p < 0.01; ***: p < 0.001; ****: p < 0.0001)

### 4.4 Surgery

Surgery was performed as previously described [25]. Anaesthetised mice were placed in a supine position, with their right leg fixed into a right-angled position. A small incision was made over the patellar tendon, and the tendon was shifted to the lateral side to expose the distal femur. A 30G needle was used to drill a hole to the intramedullary cavity and a 0.3ml insulin syringe (BD, Franklin Lakes, NJ, USA) was used to inject 10µl of cells into the intramedullary cavity. The hole was filled with bone wax (Harvard Apparatus, Holliston, MA, USA) and the skin closed with surgical clips (Michel Suture Clips, Agnthos, Lidingö, Sweden). Mice received 500µl sterile recovery saline subcutaneously.

### 4.5 Behaviour

Nociceptive behaviour – limb use and static weight bearing – was assessed in all animals in the study. Both were performed as previously described [25]. During limb use assessment, mice were observed in a transparent box for three minutes each. Mice were scored from 4 (normal gait) to 0 (complete lack of use of the ipsilateral leg and experimental endpoint). After, mice were assessed using static weight bearing in an Incapacitance Tester (version 5.2, Linton, UK). Three measurements of three seconds each was taken and the average calculated. The percentage of the total weight placed on the ipsilateral limb is reported.

Experimenters were masked/blinded to both the surgical and treatment groups until all analysis was completed.

### 4.6 Imaging

#### 4.6.1 In Vivo Imaging

All *in vivo* imaging was performed in the Lumina XR Apparatus In Vivo Imaging System (IVIS, Revitty, Belgium; exposure settings: 28 s, 35 kVp). X-ray imaging was performed as previously described, at baseline (before surgery) and at the experimental endpoint [25]. The breast cancer model contained a luciferase reporter gene that was imaged in triplicate at the experimental endpoint, as previously described [26]. Mice were injected i.p. with D-Luciferin (PerkinElmer, Inc., Shelton, CT, USA), 9 min prior to bioluminescent imaging (Binning: M(4), F/stop: 1, Exposure: 30s). An automatic region of interest was generated using a 35% peak intensity threshold and the photons/s/cm^2^ is reported.

#### 4.6.2 Micro-CT Imaging

After perfusion and prior to decalcification, the right femurs underwent micro-CT imaging according to previously published protocols [27]. Briefly, the mid-femur was scanned in a high-resolution microtomographic system (vivoCT 40, Scanco Medical AG, Switzerland). The images were scanned in high resolution and reconstructed with a voxel size of 10.5 x 10.5 x 10.5 µm^3^ (2048 x 2048 x 2048 pixels). The parameters specified in this scan included both the trabecular and cortical bone. After segmentation, the 3D microarchitectural properties of trabecular and cortical bone were quantified separately. Experimenters were masked to the surgical and treatment groups until all analysis was completed.

### 4.7 Serum Collection and Analysis

Serum was collected from mice at baseline and day 8 (4T1-Luc2 model), day 10 (RM1 model), or day 15 (NCTC2472 model) via cheek blood sampling. After behavioural tests had been performed, mice were scruffed and a sterile lancet pierced the facial vein to collect approximately 50µl of blood into a sterile 1.5ml Eppendorff tube. Samples were kept at room temperature for 1 hour and then centrifuged at 2000g for 10 min at 4°C. The serum portion was removed and placed in a new Eppendorff tube, followed by the same centrifuge procedure. The serum portion was again removed and placed in a new Eppendorff tube. Samples were stored at -80°C.

NP137 was quantified in serum by sandwich ELISA (enzyme-linked immunosorbent assay). Briefly, P96 well plates (Costar, Corning, NY, USA) were coated over night with anti-human Fab (Sigma-Aldrich, MA, USA; catalogue: I5260) at 1:2200 and blocked 1 h at room temperature with BSA (Euromedex, France; catalogue: 04-100-812-C). Samples were diluted from 1:500 to 1:2000 and loaded together with a standard of NP137. The presence of NP137 was detected with a second horse radish peroxidase (HRP)-conjugated goat anti-human IgG (Fc) (ImmunoChemistry, CA, USA; catalogue: 6291) at 1:5000, 45 min at room temperature. The signal was measured with ECL Imaging System (Thermo Scientific, MA, USA) and read on Tecan instrument Infinite M1000 PRO (Tecan, Switzerland).

Collagen biomarkers C3M (a fragment of matrix metalloprotease degraded type III collagen, nordicC3M^TM^, Nordic Bioscience, Denmark; catalogue: 1200AG) and C4M (a fragment of matrix metalloprotease degraded type IV collagen, nordicC4M^TM^, Nordic Bioscience, Denmark; catalogue: 1300AG) were measured in serum samples collected at baseline and day 15 using well-characterized and validated competitive ELISAs, according to the manufactures’ instructions (Nordic Bioscience, Denmark). In brief, the individual serum biomarkers were assessed using a 96-well streptavidin-coated plate coated with a specific biotinylated synthetic peptide dissolved in an optimized assay buffer, which incubated for 30 min at 20°C. The plate was then washed five times with washing buffer (20 mM Tris, 50 mM NaCl, pH 7.2) before adding 20 μL of either peptide calibrator or sample to the appropriate wells.

Subsequently, 100 μL of an HRP-conjugated, target-specific monoclonal antibody was added. The plate was incubated for 1 hr at 20°C, followed by five washes with washing buffer. Finally, 100 μL of tetramethylbenzidine (Kem-En-Tec, Denmark; catalogue: 438OH) was added, and the plate was incubated for 15 min at 20°C in the dark before the addition of 100 μL stopping solution (1% H_₂_SO_₄_). The optical density of each well was measured at 450 nm using 650 nm as the reference wavelength.

### 4.8 Perfusion, Fixation, and Decalcification

At the experimental endpoint, mice were anaesthetised with the ketamine/xylazine cocktail and perfuse-fixed via intracardiac perfusion. Mice were perfused with 30-40ml ice-cold PBS, followed by 30-40ml ice-cold 4% paraformaldehyde (PFA; Sigma-Aldrich, Burlington, MA, USA). The femur was removed and excess tissue removed. Samples were stored in 4% PFA overnight at 4°C, after which, bones were transferred to PBS and stored at 4°C until micro-CT imaging.

After micro-CT imaging the femurs were decalcified. Bones were placed in tissue cassettes (Tissue-Tek®, Sakura, Osaka, Japan) and placed in 10% Ethylenediaminetetraacetic acid (EDTA, ThermoFisher Scientific, Massachusetts, USA) in PBS adjusted to 7.4pH and kept at 4°C on constant rotation. EDTA was changed twice per week for 2-3 weeks until decalcification was complete, confirmed using x-ray imaging. After decalcification, bones were placed in 30% sucrose for 2-3 days and then in embedded in optimal cutting temperature media (OCT, Sakura, Osaka, Japan) and stored in -80°C until sectioning.

### 4.9 Sectioning and Immunohistochemistry

Embedded bones were placed in -25°C cryostat (CM 3050 S, Leica Microsystems, Germany) for 20-30 min to acclimatise to the temperature. The femur was cut in longitudinal sections at 25µm onto SuperFrost Plus Slides (Epredia, Runcorn, UK).

Sections were dried for 1 hour at room temperature and immediately entered IHC staining. All IHC procedures were conducted in a Sequenza Shandon Slide Rack (Epredia, Runcorn, UK). Slides were washed with 0.1M PBS, followed by blocking buffer (Normal Donkey Serum 3% (Jackson ImmunoResearch Laboratories: 017-000-121; PA, USA), 0.3% Triton X-100 (Sigma Ultra: 125K00471; Merck, Germany)) for 2 hours at room temperature. Primary calcitonin gene related peptide ((anti-rabbit; 1:3000, Sigma-Aldrich: 0000293648; Merck, Germany) in diluted blocking buffer (1:2 dilution)) were applied, then covered and left at 4°C overnight. Slides were washed three times with 0.1M PBS and then the secondary antibody ((1:600, Jackson ImmunoResearch Laboratories: 166393; MA, USA) in 1% diluted blocking buffer) were added to the slides and incubated at room temperature for 2 hours. After incubation, slides were counterstained with DAPI (4,6-diamidino-2-phenylindole; 1:20000, Sigma-Aldrich; Merck, Germany), then underwent serial dehydration in ethanol and xylene, before being coverslipped with dibutylphthalate polystyrene xylene (DPX) mounting medium (Sigma-Aldrich: BCCL3373; Merck, Germany). DPX was allowed to dry covered at room temperature. After drying, slides were stored in the dark at 4°C.

### 4.10 Western Blotting

Confluent cells were washed with cold PBS and scraped in lysis buffer (Tris 10 mM pH 7.6; sodium dodecyl sulfate (SDS) 5mM; Glycerol 10%; Triton X-100 1%, dithiothreitol 100mM). After sonication, proteins were quantified using Pierce 660nm Protein Assay Reagent (Thermo Fisher Scientific, MA, USA) and loaded on 4%-15% SDS-polyacrylamide gels (Bio-Rad, CA, USA) transferred to nitrocellulose membranes using Trans-Blot Turbo Transfer (Bio-Rad, CA, USA). Membranes were blocked for 1 hr at room temperature with 5% non-fat milk powder. Staining was performed overnight with primary antibody, anti-netrin1 antibody (Abcam, UK; catalogue: Ab126729) at 1:1000. After washing, Ntn1 stained membranes were incubated with secondary antibody, goat anti-rabbit coupled with HRP or anti-actin HRP (BioLegend, MA, USA; catalogue: 664804) at 1:5000 for 1 hr at room temperature. West Dura Chemiluminescence System (Pierce Biotechnology, MA, USA) was used to intensify the signal. Imaging was performed using ChemiDoc Touch (Bio-Rad, CA, USA).

### 4.11 Image Analysis

Nerve fibre quantification was conducted using Fiji, ImageJ 1.54f (Wayne Rasband and contributors National Institutes of Health, USA) with the NeuronJ plugin. The periosteum on both sides of the bone was analysed 2mm from the distal femur growth plate and then 2mm towards the proximal femur. The neurons within the region of interest were manually traced to determine the total length and length per periosteum area was calculated. Image acquisition and analysis was conducted by masked experimenters until all analysis was completed.

### 4.12 Statistics

All statistical analyses and graph generation was conducted in Graphpad Prism, version 11.0.2 (GraphPad Software, Inc.; NY, USA). All statistical analysis was conducted by masked experimenters. The statistical tests used and their data presentation are described in the associated figure legend.

## 5. Results

### Netrin-1 does not ameliorate cancer-induced bone pain

Six mice were excluded from the NCTC2472-vehicle group, one mouse from the RM1-vehicle group, and one mouse from the RM1-NP137, as they did not develop tumours (as visualised by x-ray imaging) during the study.

In the NCTC2472 (osteosarcoma) model, the onset of nociception development (limb use < 3; Figure 1D) was significantly delayed in the NP137-treated cancer group as compared to the vehicle-treated cancer group (p = 0.0437) (Figure 1B and 1D) and there was a significant difference (p = 0.0473) in the limb use AUC between those same groups (Figure 1B and Supplementary Figure 1C).; There was no significant difference between the NP137- and vehicle-treated cancer groups for weight bearing (Figure 1C and Supplementary Figure 1D). In the 4T1-Luc2-inoculated (breast cancer) study, there was no significant difference between the vehicle- and NP137-treated groups in both the limb use (Figure 1E; Supplementary Figure 1I) and weight bearing (Figure 1F; Supplementary Figure 1J) tests, and there was no delay in nociception onset (Figure 1G). In the RM1-inoculated model (prostate cancer), a similar trend was observed with no significant difference between the NP137- and vehicle-treated cancer groups in both limb use and weight bearing (Figure H-J; Supplementary Figure 1O-P). In all models, across the full experimental timeline, overall survival was not significantly improved (Supplementary Figure 1A, G, M) and there was no significant difference in weight between the groups (Supplementary Figure 1B, H, N). In all models, the sham groups did not exhibit nociceptive behaviour or tumour development and there was no significant difference between the NP137- and vehicle-treated sham groups (Figure 1 and Supplementary Figure 1).

### Bone destruction accompanies tumour development in all models

Tumour development in the bone results in trabecular and cortical bone destruction through osteoclastic bone resorption (NCTC2472 and 4T1-Luc2 models) or osteoblastic/osteolytic abnormal development (RM1 model). MicroCT analysis was performed to measure bone structure parameters indicative of bone destruction (Supplementary Table 1).

In the NCTC2472-inoculated and 4T1-Luc2 model, the bone volume-to-total volume ratio, trabecular mean density, trabecular thickness, and trabecular separation were all significantly deteriorated compared to their sham-treatment controls and there was no significant difference between treatment groups (Figure 2A-H). The 4T1-Luc2 contains a luciferase reporter gene which was used to assess tumour growth in the model. There was no significant difference between the vehicle- and NP137-treated groups (Figure 2I).

**Figure 2:**
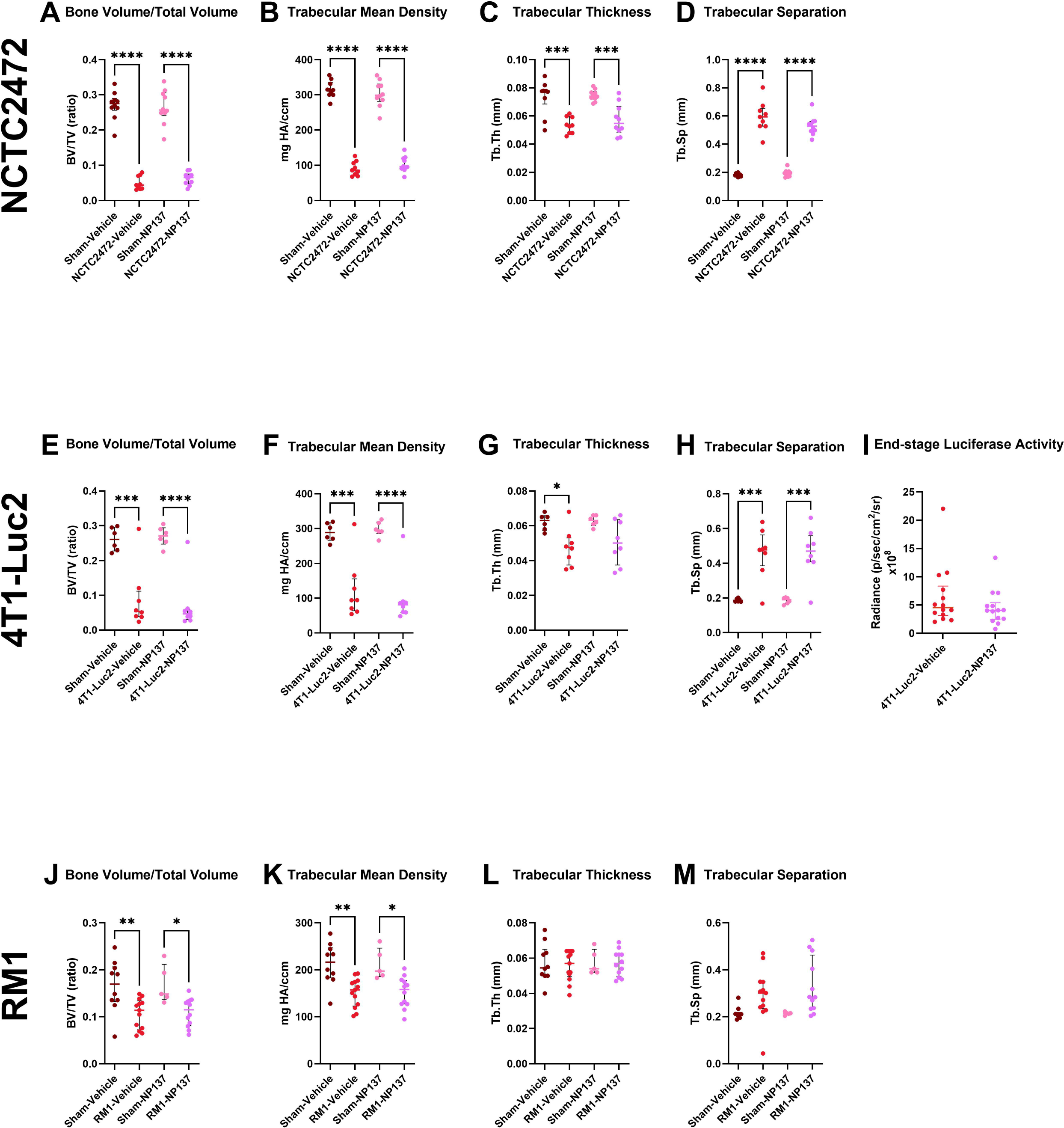
Micro-computed tomography analysis of ipsilateral femurs, and luciferase imaging of the 4T1-Luc2 model. (A-D) Micro-CT analysis of ipsilateral femurs taken from the NCTC2472 model; (E-H) micro-CT analysis of ipsilateral femurs taken from the 4T1-Luc2 model; (I) luciferase activity of inoculated 4T1-Luc2 cells taken just prior to experimental endpoint; (J-M) micro-CT analysis of ipsilateral femurs taken from the RM1 model. (A, E, J) Bone volume-to-total volume ratio; (B, F, K) mean density of the trabecular bone; (C, G, L) trabecular thickness; (D, H, M) trabecular separation. All graphs display the median±IQR and all statistical analyses are one-way ANOVA with Tukey test. (BV: bone volume; TV: total volume; Tb: trabecular; Th: thickness; Sp: separation; mg HA/ccm: milligrams of Hydroxyapatite per cubic centimetre; mm: millimetre; p/sec/cm2/sr: photons per second per square centimeter per steradian; *: p < 0.05; **: p < 0.01; ***: p < 0.001; ****: p < 0.0001)

In the RM1 model, the bone volume-to-total volume ratio and trabecular mean density were significantly deteriorated compared to their sham-treatment controls and none of the parameters showed significant differences between the NP137- and vehicle treated cancer groups (Figure 2J-M). In all models, vehicle- and NP137-treated mice in the sham groups did not exhibit significantly different change for any measurements (Figure 2).

### Co-administration of NP137 and etanercept does not produce anti-nociception

Recent studies have indicated that co-inhibition of both nerve growth factor (NGF) and tumour necrosis factor-alpha (TNF-α) results in significant nociception reduction. The reduction is similar to or better than treatment with the drugs individually [10], [28]. Therefore, we decided to test whether co-administration of NP137 with etanercept (TNF-α inhibitor) could improve analgesic outcomes in the RM1 model. In the limb use and weight bearing tests, there was no significant difference between the any of the RM1-inoculated groups during the course of the study (Figure 3A-D). There was no significant difference in survival or time of nociceptive onset between any of the RM1-inoculated groups (Figure 3E-F) and the AUC for limb use and weight bearing did not show significant differences between any of cancer groups (Figure 3G-H). During the course of the study, we noted that lung metastases occurred in approximately half of the mice (Figure 3I). In the vehicle only-treated group, most of the mice developed lung metastases (7 mice), half of which reached the humane endpoint prior to the experimental endpoint (lack of use of the ipsilateral limb due to nociception development) and were excluded from the study. This resulted in an underpowered group. However, mice treated with etanercept and NP137 developed far fewer metastases (2-3 mice per group), indicating that these drugs might protect against metastatic progression in the RM1 model.

**Figure 3:**
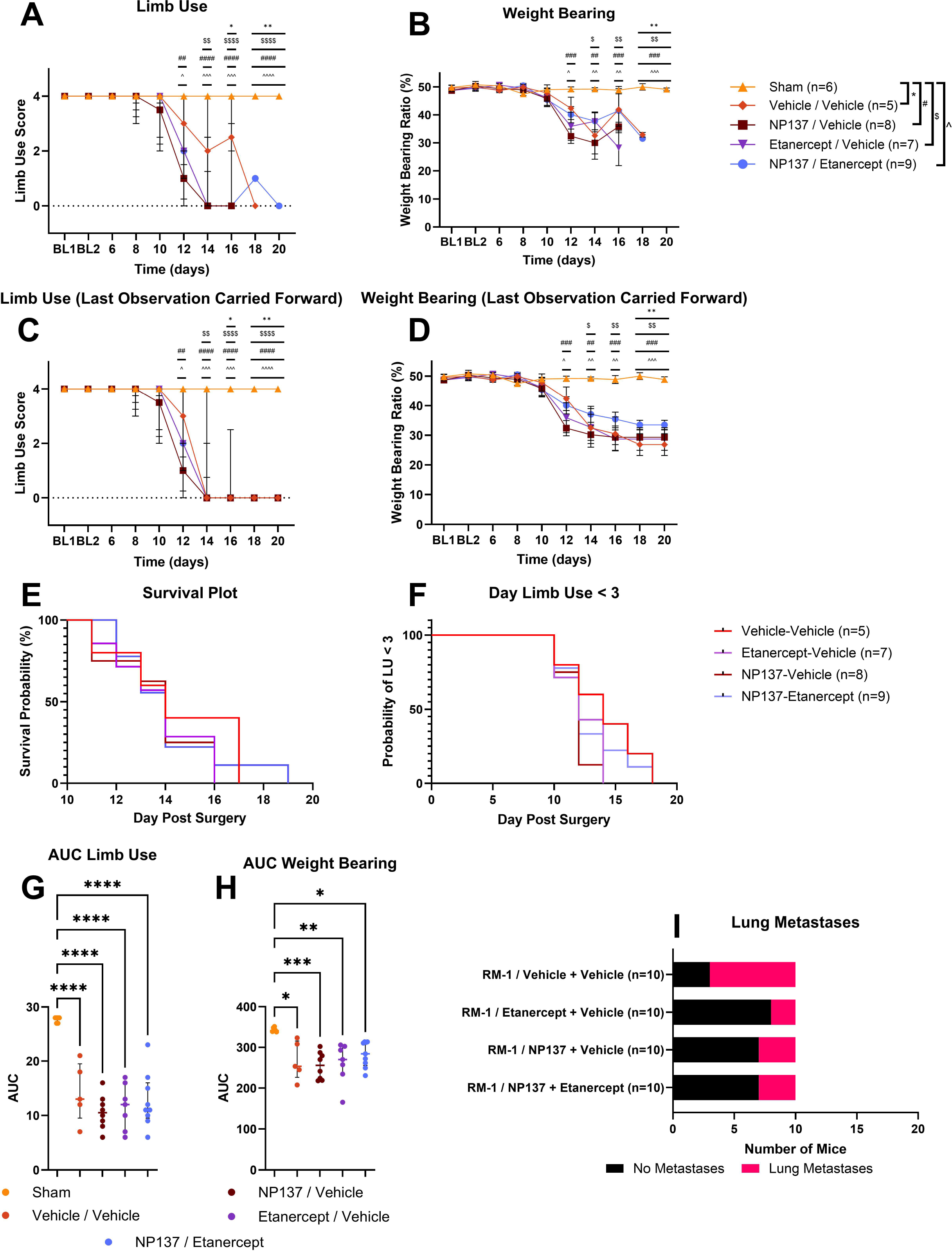
Combination treatment behaviour outcomes. (A) Limb use behaviour represented as median±IQR (two-way ANOVA with Dunnett Test); (B) weight bearing behaviour represented as mean±SEM (two-way ANOVA with Dunnett Test); (C) limb use behaviour showing last observation carried forward for statistical analysis (two-way ANOVA with Dunnett Test); (D) weight bearing behaviour showing last observation carried forward for statistical analysis (two-way ANOVA with Dunnett Test); (E) survival plot for RM1-inoculated mice (log-rank survival test); (F) survival plot displaying day at which a limb use score below 3 was achieved (log-rank survival test); (G) limb use behaviour AUC from day 6 showing median±IQR (one-way ANOVA with Tukey Test); (H) weight bearing behaviour AUC from day 6 showing median±IQR (one-way ANOVA with Tukey Test); (I) proportion of groups developing lung metastases. (BL: baseline; n: number; AUC: area under the curve; IQR: interquartile range; SEM: standard error of the mean; ANOVA; analysis of variance; *: p < 0.05; **: p < 0.01; ***: p < 0.001; ****: p < 0.0001)

### Cancer development does not deplete circulating NP137

Netrin-1 is primarily known for its role in nervous system development during foetal development, but more recently netrin-1 has been shown to be produced by cancer cells and osteoclasts [20], [24]. Western blotting was performed on the three cell lines used to determine whether they produced netrin-1. The 4T1-Luc2 and RM1 cells both produced detectable netrin-1, but the NCTC2472 cell line did not (Figure 4A). Previous studies on the analgesic efficacy of NP137 in cancer-induced bone pain indicated that circulating NP137 was significantly depleted in the cancer-inoculated group compared to the sham-operated group [23]. To determine whether the treatment strategy (loading dose with continued dosing) sustained circulating NP137, an ELISA was conducted to determine NP137 in serum. There was no significant difference between the sham and cancer NP137-treated groups, indicating that the dosing strategy maintained sustained NP137 serum concentration (Figure 4B).

**Figure 4:**
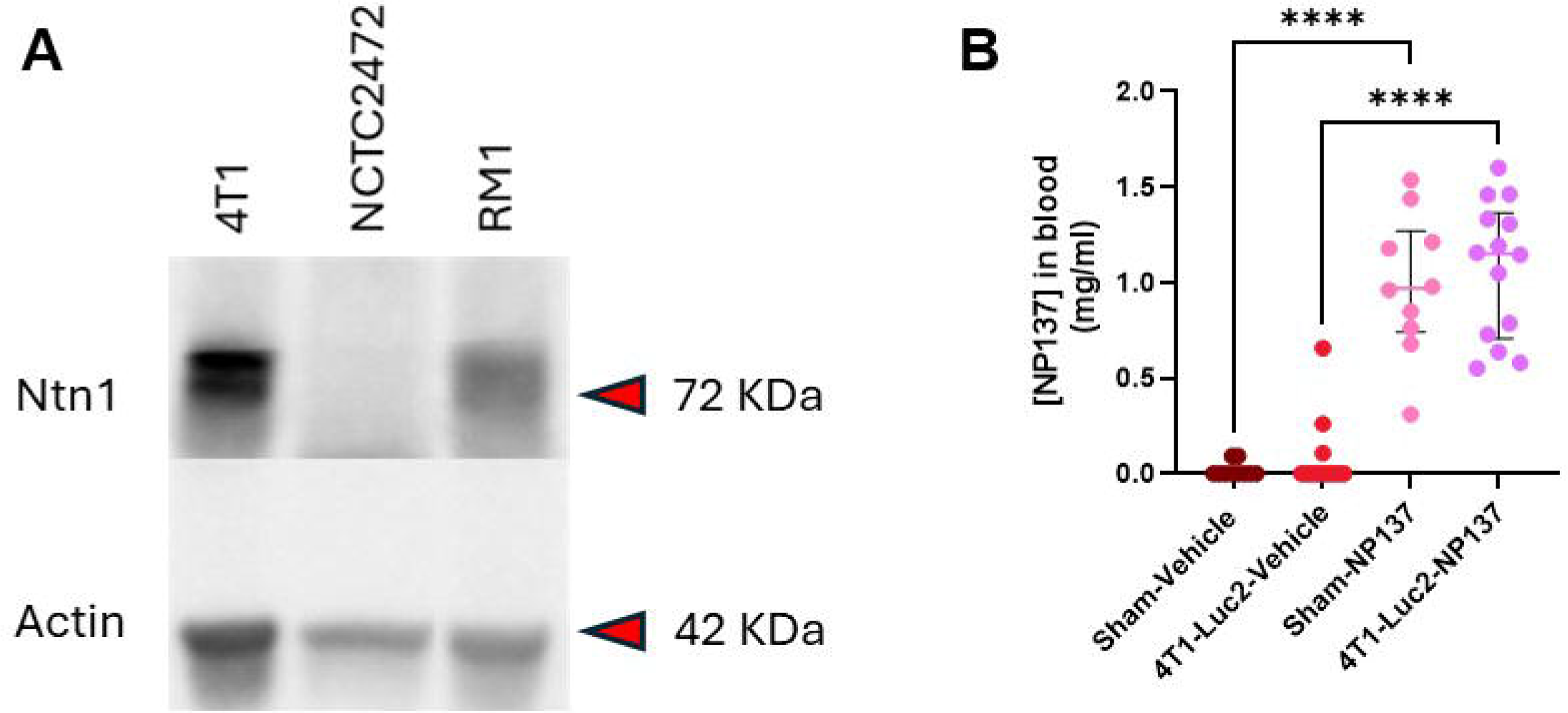
Netrin-1 and NP137 exposure assessments. (A) Western blot displaying netrin-1 (Ntn1) presence in the three cancer cell lines. Beta-actin (Actin) was used as a housekeeping control; (B) serum concentration of NP137 taken at experimental endpoint. Analysis conducted with one-way ANOVA with Tukey Test and presented as median±IQR. (KDa: kiloDalton; mg/ml: milligrams/millilitres; n: number; ****: p < 0.0001).

Analysis of calcitonin gene related peptide nerve fibres in the bone showed no difference between any of the groups in any of the models (Supplementary Figure 2A-C). Serum was also used to assess collagen markers, which may indicate tissue destruction and remodelling (e.g. bone basement membrane, synovial tissue, etc.). The NCTC2472 and RM1 models did not demonstrate serum C3M or C4M differences between sham and cancer groups at baseline and day 15, and there were no concentration changes after NP137 treatment (Supplementary Figure 2D-G).

## 6. Discussion

This study aimed to determine the analgesic efficacy of NP137 treatment (netrin-1 inhibition) in three models of cancer-induced bone pain – osteosarcoma, metastatic breast cancer, and metastatic prostate cancer.

NP137 produced a delay in nociception in the osteosarcoma model, but no significant effects in the breast and prostate cancer models. However, a minor transient effect was observed in all three models in the early- to mid-stages of nociception development. This supports previous studies on NP137 in multiple myeloma-induced pain, in which a transient analgesic effect was observed following NP137 treatment [23]. It is interesting that the same trends were broadly observed in four different models of CIBP. This indicates that netrin-1 might have a biological effect in CIBP in the early-stages of disease progression. We are speculating that nerve sprouting occurs in this stage of the disease having recently found increased periosteal nerve density in the NCTC2472 model at the mid-stage [25]. The nerve sprouting may therefore also be transient, as this study demonstrated that there was no change in nerve density when observed at the late stage.

Previous studies have shown that NP137 has the ability to act as an analgesic [21], prompting the question as to why NP137 fails to reduce nociception in these models? We have demonstrated that there is sufficient NP137 in the serum, suggesting that the lack of nociceptive effect is not due to poor exposure.

CIBP is the summation of many nociceptive and neuropathic mechanisms that affect both the peripheral and central nervous systems. In the peripheral nervous system, cells within the cancer-bone environment produce inflammatory mediators, tissue damage occurs, and nerves are damaged, and in the central nervous system, central sensitisation occurs [1]. This indicates that the nociceptive mechanisms may be complex, controlled by various molecular factors, vary between models, and may indeed occur in a stage-specific manner.

In this study, we observed variability between the models, in which the NCTC2472 model produced a significant transient effect despite the cells not producing netrin-1. Previous studies have shown that osteoclasts produce netrin-1 [20], which may suggest why NP137 has an anti-nociceptive effect and why it is different from the 4T1-Luc2 and RM1 models. Further studies would determine the role of netrin-1 in non-cancer cells of the bone environment.

As such, even if netrin-1 is a pronociceptive mechanism in CIBP, targeting a single mechanism is unlikely to be efficacious when there are many other mechanisms activating nociceptive pathways. CIBP treatment is likely to benefit from an individualised and multi-drug approach targeting multiple nociceptive pathways. Netrin-1 inhibition may prove to be efficacious using such an approach in early cancer development; however, this study has shown that it is unlikely to be efficacious on its own.

## 7. Conclusions

In this study, netrin-1 inhibition via NP137 treatment did not ameliorate cancer-induced bone pain in models of osteosarcoma, metastatic breast cancer, or metastatic prostate cancer. A consistent transient effect has been observed in four models [23] of cancer-induced bone pain in early stages, but ultimately, nociception and disease progression remain unaffected. This study has shown that NP137 cannot provide analgesia for CIBP; however, if suitable co-compounds are identified, NP137 may be a beneficial adjuvant therapy for CIBP in early stages.

## 8. Figure Legands

**Supplementary Figure 1:** Supplementary behavioural and in vivo observations. (A-F) NCTC2472 model results; (G-L) 4T1-Luc2 model results; (M-R) RM1 model results; (A, G, H) survival plots of cancer inoculated groups (analysed with log-rank test); (B, H, N) mouse weights over the course of the study (mean±SEM; analysed using two-way ANOVA with Tukey Test); (C, I, O) limb use AUC (presented as median±IQR; analysed using one-way ANOVA with Tukey Test); (D, J, P) weight bearing AUC (presented as median±IQR; analysed using one-way ANOVA with Tukey Test); (E, K, Q) limb use behaviour over time with last observation carried forward (presented as median±IQR; analysed using two-way ANOVA with Tukey Test); (F, L, R) weight bearing behaviour over time with last observation carried forward (presented as mean±SEM; analysed using two-way ANOVA with Tukey Test).

(BL: baseline; n: number; AUC: area under the curve; IQR: interquartile range; SEM: standard error of the mean; ANOVA; analysis of variance; *: p < 0.05; **: p < 0.01; ***: p < 0.001; ****: p < 0.0001)

**Supplementary Figure 2:** Nerve fibre density analysis and serum markers of collagen degradation. (A-C) CGRP nerve fibre density in ipsilateral femurs measured using NeuronJ (presented as median±IQR; analysed using two-way ANOVA with Tukey Test); (D) C3M serum concentration from NCTC2472 model at baseline and day 15 (presented as median±IQR; analysed using two-way ANOVA with Tukey Test); (E) C4M serum concentration from NCTC2472 model at baseline and day 15 (presented as median±IQR; analysed using two-way ANOVA with Tukey Test); (F) C3M serum concentration from RM1 model at baseline and day 10 (presented as median±IQR; analysed using two-way ANOVA with Tukey Test); (G) C3M serum concentration from RM1 model at baseline and day 10 (presented as median±IQR; analysed using two-way ANOVA with Tukey Test).

(BL: baseline; n: number; IQR: interquartile range; D: day; CGRP: calcitonin gene related peptide; µm: micrometre; C3M: type III collagen fragment; C4M: type IV collagen fragment; ANOVA; analysis of variance; *: p < 0.05; **: p < 0.01; ***: p < 0.001; ****: p < 0.0001)

**Supplementary Table 1:** 3-D microarchitectural properties of trabecular and cortical bones, presented as mean ± standard deviation. Italics represent measurements demonstrating significant difference from the corresponding sham-vehicle treated group (surrounded by dark border). All statistical analyses are one-way ANOVA with Tukey test.

(SD: standard deviation; BS: bone surface; BV: bone volume; TV: total volume; CD: connectivity density; SMI: structure model index; Tb: trabecular bone; Ct; cortical bone; Th: thickness; Sp: separation: N: number; mm: millimetre; mg HA/ccm: milligrams of Hydroxyapatite per cubic centimetre; *: p<0.05; **: p<0.01; ***: p<0.001; ****: p<0.0001)

## 9. Acknowledgements

We would like to thank and acknowledge the support and assistance from Netris Pharma, particularly Benjamin Ducarouge and Patrick Mehlen. Netris Pharma gifted NP137, advised on drug administration, and performed NP137 and netrin-1 detection. We acknowledge the Core Facility for Integrated BioImaging, Faculty of Health and Medical Sciences, University of Copenhagen. We also acknowledge the technical support and assistance provided by Camilla Skånstrøm Dall, Mathilde Caldara, Helena Skourup Larsen, Natascha Synnøve Olsen, and Sascha A. Petersen.

## 10. CRediT Author Statement

Chelsea Hopkins: Conceptualization, Methodology, Software, Validation, Formal analysis, Investigation, Data Curation, Writing - Original Draft, Writing - Review & Editing, Visualization, Supervision, Project administration

Mie Brandt Lassen: Methodology, Formal analysis, Investigation, Data Curation, Writing - Review & Editing, Visualization

Louise Ploug Hansen: Methodology, Formal analysis, Investigation, Data Curation, Writing - Review & Editing, Visualization

Yingyu Tang: Formal analysis, Investigation, Data Curation, Writing - Review & Editing, Visualization

Edward Ciputra: Formal analysis, Investigation, Data Curation, Writing - Review & Editing, Visualization

Theis Lund Jørgensen: Formal analysis, Investigation, Data Curation, Writing - Review & Editing, Visualization

Mads Haaber Christensen: Formal analysis, Investigation, Data Curation, Writing - Review & Editing, Visualization

Cecilie Louise Pedersen: Methodology, Formal analysis, Investigation, Data Curation, Writing - Review & Editing, Visualization

Camilla I Svensson: Conceptualization, Methodology, Writing - Review & Editing, Funding acquisition

Ming Ding: Formal analysis, Writing - Review & Editing, Visualization, Supervision

Rasmus S Pedersen: Investigation, Data Curation, Writing - Review & Editing

Nicholas Willumsen: Conceptualization, Methodology, Formal analysis, Resources, Writing - Review & Editing, Supervision, Funding acquisition

Anne-Marie Heegaard: Conceptualization, Methodology, Software, Validation, Formal analysis, Resources, Data Curation, Writing - Original Draft, Writing - Review & Editing, Supervision, Project administration, Funding acquisition

## References

[1] S. Falk and A. H. Dickenson, ‘Pain and Nociception: Mechanisms of Cancer-Induced Bone Pain’, Journal of Clinical Oncology, Accessed: Apr. 15, 2026. [Online]. Available: https://ascopubs.org/doi/10.1200/JCO.2013.51.7219

[2] H. Smith, ‘A Comprehensive Review of Rapid-Onset Opioids for Breakthrough Pain’, CNS Drugs, vol. 26, no. 6, pp. 509–535, Jun. 2012, doi: 10.2165/11630580-000000000-00000.

[3] A. Colosia et al., ‘The Burden of Metastatic Cancer-Induced Bone Pain: A Narrative Review’, JPR, vol. 15, pp. 3399–3412, Oct. 2022, doi: 10.2147/JPR.S371337.

[4] A. A. Anekar, J. M. Hendrix, and M. Cascella, ‘WHO Analgesic Ladder’, in StatPearls, Treasure Island (FL): StatPearls Publishing, 2026. Accessed: Apr. 15, 2026. [Online]. Available: http://www.ncbi.nlm.nih.gov/books/NBK554435/

[5] J. Poço Gonçalves, D. Veiga, and A. Araújo, ‘Chronic pain, functionality and quality of life in cancer survivors’, British Journal of Pain, vol. 15, no. 4, pp. 401–410, Nov. 2021, doi: 10.1177/2049463720972730.

[6] C. Pérez et al., ‘Pain in Long-Term Cancer Survivors: Prevalence and Impact in a Cohort Composed Mostly of Breast Cancer Survivors’, Cancers, vol. 16, no. 8, p. 1581, Jan. 2024, doi: 10.3390/cancers16081581.

[7] R. Zajączkowska, M. Kocot-Kępska, W. Leppert, and J. Wordliczek, ‘Bone Pain in Cancer Patients: Mechanisms and Current Treatment’, International Journal of Molecular Sciences, vol. 20, no. 23, p. 6047, Jan. 2019, doi: 10.3390/ijms20236047.

[8] J. R. Ghilardi et al., ‘Administration of a Tropomyosin Receptor Kinase Inhibitor Attenuates Sarcoma-Induced Nerve Sprouting, Neuroma Formation and Bone Cancer Pain’, Mol Pain, vol. 6, pp. 1744–8069-6–87, Jan. 2010, doi: 10.1186/1744-8069-6-87.

[9] J. M. Jimenez-Andrade et al., ‘Pathological Sprouting of Adult Nociceptors in Chronic Prostate Cancer-Induced Bone Pain’, J Neurosci, vol. 30, no. 44, pp. 14649–14656, Nov. 2010, doi: 10.1523/JNEUROSCI.3300-10.2010.

[10] R. Haroun et al., ‘Novel therapies for cancer-induced bone pain’, Neurobiology of Pain, vol. 16, p. 100167, Jul. 2024, doi: 10.1016/j.ynpai.2024.100167.

[11] J. A. Carrino et al., ‘Characterization of adverse joint outcomes in patients with osteoarthritis treated with subcutaneous tanezumab’, Osteoarthritis and Cartilage, vol. 31, no. 12, pp. 1612–1626, Dec. 2023, doi: 10.1016/j.joca.2023.08.010.

[12] E. Y. van Battum, M. H. van den Munkhof, and R. J. Pasterkamp, ‘Novel insights into the regulation of neuron migration by axon guidance proteins’, Current Opinion in Neurobiology, vol. 92, p. 103012, Jun. 2025, doi: 10.1016/j.conb.2025.103012.

[13] J. Li, G. Wang, Y. Weng, M. Ding, and W. Yu, ‘Netrin-1 contributes to peripheral nerve injury induced neuropathic pain via regulating phosphatidylinositol 4-kinase IIa in the spinal cord dorsal horn in mice’, Neuroscience Letters, vol. 735, p. 135161, Sep. 2020, doi: 10.1016/j.neulet.2020.135161.

[14] C.-H. Wu et al., ‘Netrin-1 Contributes to Myelinated Afferent Fiber Sprouting and Neuropathic Pain’, Mol Neurobiol, vol. 53, no. 8, pp. 5640–5651, Oct. 2016, doi: 10.1007/s12035-015-9482-x.

[15] J. T.-C. Chen, L. Schmidt, C. Schürger, M. K. Hankir, S. M. Krug, and H. L. Rittner, ‘Netrin-1 as a Multitarget Barrier Stabilizer in the Peripheral Nerve after Injury’, International Journal of Molecular Sciences, vol. 22, no. 18, p. 10090, Jan. 2021, doi: 10.3390/ijms221810090.

[16] A. Köhle et al., ‘Serum netrin-1 and netrin receptor levels in fibromyalgia and osteoarthritis’, Turk J Phys Med Rehabil, vol. 68, no. 2, pp. 238–245, Jun. 2022, doi: 10.5606/tftrd.2022.8114.

[17] B. Zheng et al., ‘Netrin-1 mediates nerve innervation and angiogenesis leading to discogenic pain’, J Orthop Translat, vol. 39, pp. 21–33, Dec. 2022, doi: 10.1016/j.jot.2022.11.006.

[18] S. Ding et al., ‘Macrophage-derived netrin-1 contributes to endometriosis-associated pain’, Annals of Translational Medicine, vol. 9, no. 1, pp. 29–29, Jan. 2021, doi: 10.21037/atm-20-2161.

[19] H. Wei, L. Ailanen, M. Morales, A. Koivisto, and A. Pertovaara, ‘Spinal TRPA1 Contributes to the Mechanical Hypersensitivity Effect Induced by Netrin-1’, Int J Mol Sci, vol. 23, no. 12, p. 6629, Jun. 2022, doi: 10.3390/ijms23126629.

[20] S. Zhu et al., ‘Subchondral bone osteoclasts induce sensory innervation and osteoarthritis pain’, J Clin Invest, vol. 129, no. 3, pp. 1076–1093, Mar. 2019, doi: 10.1172/JCI121561.

[21] R. Rudjito, et al., ‘Bone innervation and vascularization regulated by osteoclasts contribute to refractive pain-related behavior in the collagen antibody-induced arthritis model’, Apr. 20, 2021, bioRxiv. doi: 10.1101/2021.04.19.440384.

[22] Z. Gong et al., ‘Netrin-1 Role in Nociceptive Neuron Sprouting through Activation of DCC Signaling in a Rat Model of Bone Cancer Pain’, J. Integr. Neurosci., vol. 23, no. 3, Feb. 2024, doi: 10.31083/j.jin2303047.

[23] M. Diaz-delCastillo et al., ‘Metastatic Infiltration of Nervous Tissue and Periosteal Nerve Sprouting in Multiple Myeloma-Induced Bone Pain in Mice and Human’, J. Neurosci., vol. 43, no. 29, pp. 5414–5430, Jul. 2023, doi: 10.1523/JNEUROSCI.0404-23.2023.

[24] P. A. Cassier et al., ‘Netrin-1 blockade inhibits tumour growth and EMT features in endometrial cancer’, Nature, vol. 620, no. 7973, pp. 409–416, Aug. 2023, doi: 10.1038/s41586-023-06367-z.

[25] J.-P. Fuller-Jackson, C. Hopkins, J. Thai, M. B. Lassen, A.-M. Heegaard, and J. Ivanusic, ‘Three-Dimensional Analysis of the Effect of Osteosarcoma on Sensory Nerves Innervating the Femur in a Murine Model of Osteosarcoma-Induced Bone Pain’, Cancers (Basel), vol. 17, no. 21, p. 3533, Oct. 2025, doi: 10.3390/cancers17213533.

[26] C. Hopkins, I. B. Kanneworff, B. R. Kornum, and A.-M. Heegaard, ‘Wheel Running in Digital Ventilated Cages® Is Impaired in a Model of Cancer-induced Bone Pain’, In Vivo, vol. 39, no. 6, pp. 3205–3215, Nov. 2025, doi: 10.21873/invivo.14120.

[27] C. H. Dreyer, M. Rasmussen, R. H. Pedersen, S. Overgaard, and M. Ding, ‘Comparisons of Efficacy between Autograft and Allograft on Defect Repair In Vivo in Normal and Osteoporotic Rats’, BioMed Research International, vol. 2020, no. 1, p. 9358989, 2020, doi: 10.1155/2020/9358989.

[28] K. M. Scott et al., ‘A Randomized, Double-Blind, Placebo-Controlled, Dose-Response Phase 2a Study of the Efficacy and Safety of a Bispecific Fusion Protein (MEDI7352) Targeting NGF and TNFα in Patients with Painful Diabetic Neuropathy’, Nov. 30, 2025, medRxiv. doi: 10.1101/2025.11.27.25341159.

[29] K. G. Halvorson, M. A. Sevcik, J. R. Ghilardi, T. J. Rosol, and P. W. Mantyh, ‘Similarities and Differences in Tumor Growth, Skeletal Remodeling and Pain in an Osteolytic and Osteoblastic Model of Bone Cancer’, The Clinical Journal of Pain, vol. 22, no. 7, p. 587, Sep. 2006, doi: 10.1097/01.ajp.0000210902.67849.e6.

