## Supplementary figures and images for "Netrin-1 inhibition does not attenuate cancer-induced bone pain in three translational models"

### Supplementary Figure 1

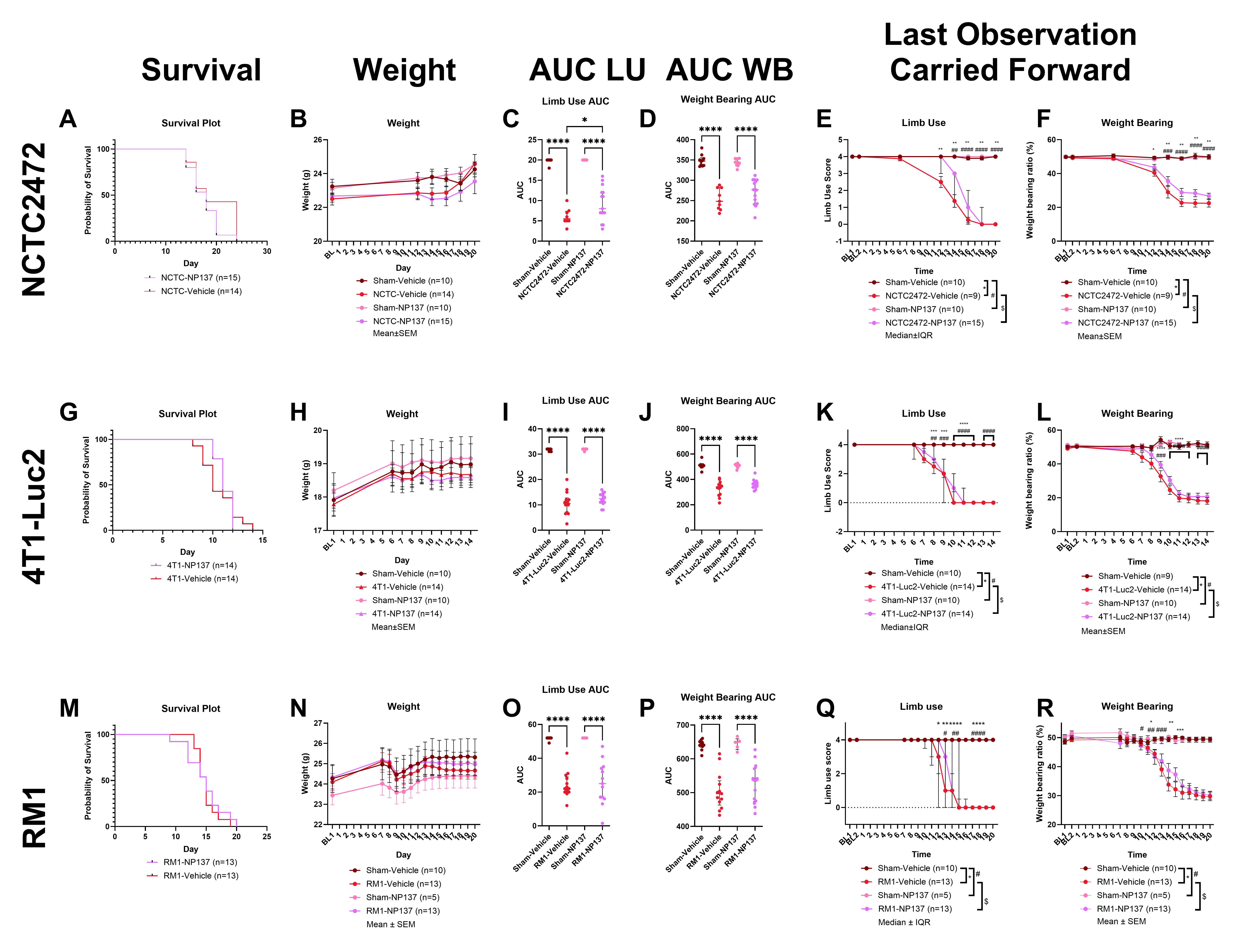

### Supplementary Figure 2

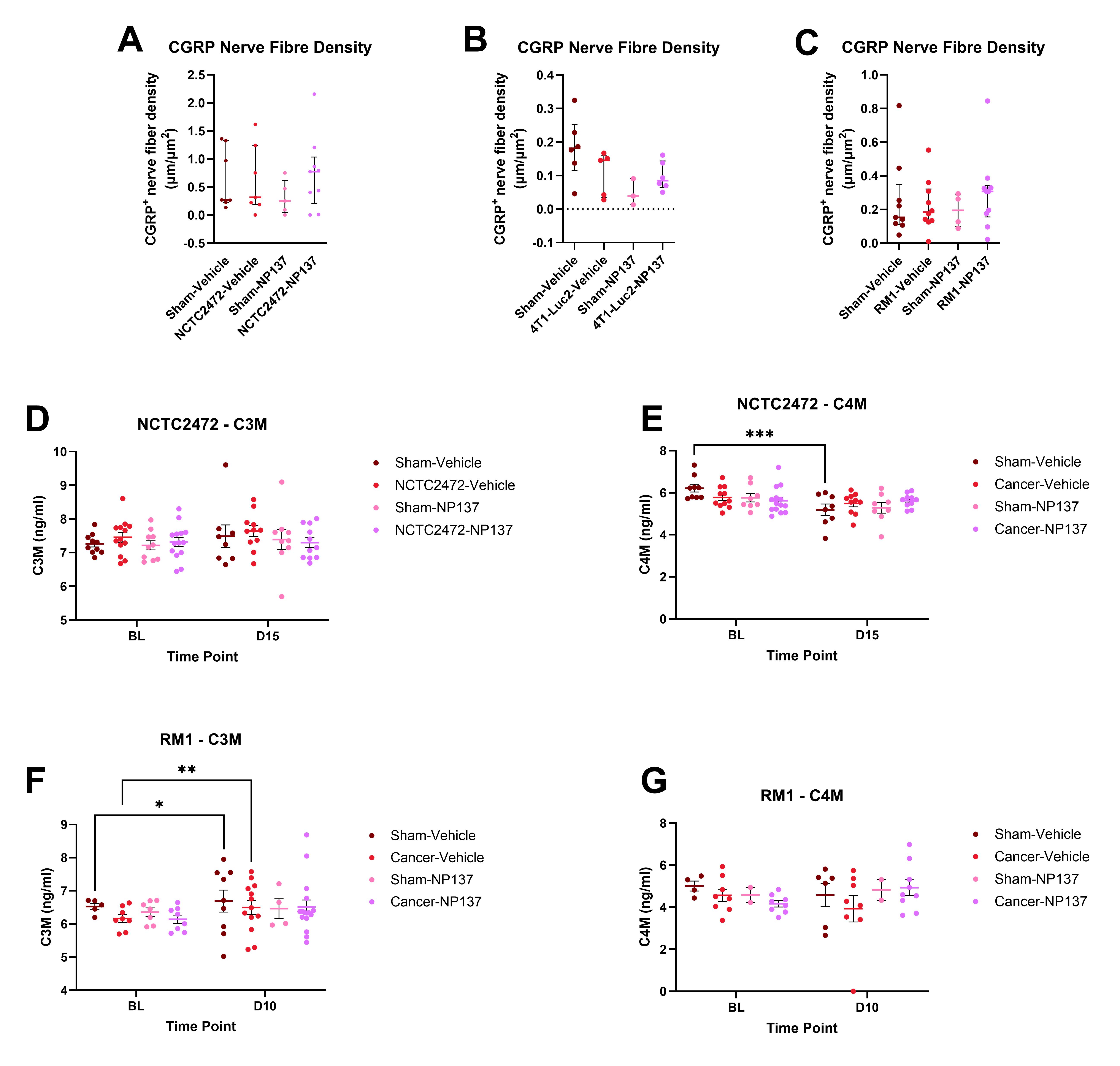
