## Supplementary Table 1 for "Netrin-1 inhibition does not attenuate cancer-induced bone pain in three translational models"

Supplementary Table 1: 3-D microarchitectural properties of trabecular and cortical bones, presented as mean ± standard deviation.

|  |  |  | **Sham-Vehicle (mean±SD)** | **Cancer-Vehicle (mean±SD)** | **Sham-NP137 (mean±SD)** | **Cancer-NP137 (mean±SD)** |
| --- | --- | --- | --- | --- | --- | --- |
| **Osteosarcoma Cancer (NCTC2472 Inoculation)** | **Trabecular bone** | **BV/TV** | **0.270*±*0.040** | ***0.050±0.019******* | **0.265*±*0.049** | ***0.063±0.018******* |
|  |  | BS/BV (mm^-1^) | 35.320±2.158 | *48.320±4.475***** | 36.930±2.961 | *48.220±5.607***** |
|  |  | BS/TV (mm^-1^) | 9.962*±*0.518 | *2.370±0.728***** | 9.661*±*1.141 | *2.961±0.748***** |
|  |  | CD  (mm^-3^) | 183.200*±*14.680 | *17.360±6.398***** | 172.900*±*17.550 | *33.910±23.100***** |
|  |  | SMI | 1.485*±*0.159 | *2.291±0.164***** | 1.509*±*0.301 | *2.151±0.331***** |
|  |  | Tb.Th (mm) | 0.074*±*0.012 | *0.054±0.006**** | 0.075*±*0.004 | *0.057±0.011*** |
|  |  | Tb.Sp (mm) | 0.183*±*0.010 | *0.594±0.109***** | 0.194*±*0.025 | *0.530±0.068***** |
|  |  | Tb.N | 4.914*±*0.223 | *1.813±0.308***** | 4.748*±*0.436 | *1.935±0.228***** |
|  |  | Apparent Density (mg HA/ccm) | 316.100*±*23.660 | *90.860±20.490***** | 303.000*±*36.710 | *103.000±21.950***** |
|  |  | Material Density (mg HA/ccm) | 849.200*±*18.310 | *793.500±31.170***** | 845.200*±*13.670 | *772.300±27.420***** |
|  | **Cortical Bone** | BS/BV (mm^-1^) | 17.460*±*6.363 | *25.310±1.769*** | 14.440*±*0.985 | *23.970±5.452** |
|  |  | BS/TV (mm^-1^) | 8.611*±*0.573 | *5.948±0.520***** | 7.978*±*0.685 | *5.868±0.872***** |
|  |  | Porosity (%) | 0.478*±*0.090 | *0.7638±0.028***** | 0.448*±*0.026 | *0.739±0.080***** |
|  |  | Ct.Th (mm) | 0.150*±*0.027 | *0.113±0.009** | 0.173*±*0.018 | 0.123*±*0.035 |
|  |  | Ct.Sp (mm) | 0.175*±*0.012 | *0.393±0.052***** | 0.185*±*0.024 | *0.414±0.047***** |
|  |  | Apparent Density (mg HA/ccm) | 521.500*±*79.760 | *239.100±27.150***** | 559.300*±*21.930 | *263.200±81.590***** |
|  |  | Material Density (mg HA/ccm) | 955.200*±*52.930 | *815.400±37.400***** | 987.900*±*27.180 | *826.300±66.180***** |
| **Breast Cancer (4T1-Luc2 Inoculation)** | **Trabecular bone** | **BV/TV** | **0.261±0.034** | ***0.088±0.087****** | **0.269±0.028** | ***0.070±0.075******* |
|  |  | BS/BV (mm^-1^) | 37.840±7.506 | 52.130±7.159 | 34.240±16.560 | 43.350±19.220 |
|  |  | BS/TV (mm^-1^) | 10.020±2.160 | *4.127±3.250** | 9.308±4.589 | *3.006±3.615*** |
|  |  | CD  (mm^-3^) | 276.900±23.280 | *80.200±90.770**** | 287.900±53.050 | *64.460±106.800**** |
|  |  | SMI | 1.131±0.241 | *1.938±0.424*** | 1.077±0.183 | *2.173±0.499**** |
|  |  | Tb.Th (mm) | 0.062*±*0.005 | *0.048±0.011** | 0.063*±*0.003 | 0.051*±*0.013 |
|  |  | Tb.Sp (mm) | 0.188*±*0.010 | *0.457±0.144**** | 0.185*±*0.018 | *0.460±0.144**** |
|  |  | Tb.N | 5.270*±*0.274 | *2.555±1.328**** | 5.395*±*0.499 | *2.585±1.234**** |
|  |  | Apparent Density (mg HA/ccm) | 289.300*±*26.920 | *122.700±84.940**** | 298.000*±*21.220 | *98.290±74.330***** |
|  |  | Material Density (mg HA/ccm) | 836.700±9.867 | 815.700±35.080 | 854.000±30.250 | 822.000±43.000 |
|  | **Cortical Bone** | BS/BV (mm^-1^) | 0.644±0.158 | 1.379±1.612 | 0.657±0.161 | 1.355±0.626 |
|  |  | BS/TV (mm^-1^) | 0.610±0.152 | 1.223±1.309 | 0.620±0.155 | 1.252±0.567 |
|  |  | Porosity (%) | 0.054±0.008 | 0.073±0.040 | 0.057±0.009 | 0.070±0.018 |
|  |  | Ct.Th (mm) | 0.218±0.022 | 0.200±0.020 | 0.211±0.014 | 0.214±0.022 |
|  |  | Ct.Sp (mm) | 0.012±0.004 | 0.028±0.029 | 0.018±0.009 | 0.033±0.025 |
|  |  | Apparent Density (mg HA/ccm) | 1166.0±27.460 | 1142.0±45.900 | 1161.0±30.440 | 1146.0±34.510 |
|  |  | Material Density (mg HA/ccm) | 1261.0±23.440 | 1256.0±20.250 | 1259.0±28.250 | 1257.0±24.600 |
| **Prostate Cancer (RM1 Inoculation)** | **Trabecular bone** | **BV/TV** | **0.166*±*0.055** | ***0.107±0.032***** | **0.169*±*0.041** | ***0.110±0.030**** |
|  |  | BS/BV (mm^-1^) | 48.580*±*8.332 | 50.260*±*6.993 | 47.630*±*5.866 | 48.440*±*5.941 |
|  |  | BS/TV (mm^-1^) | 7.633*±*1.620 | *5.206±1.183**** | 7.870*±*0.877 | *5.270±1.406*** |
|  |  | CD  (mm^-3^) | 142.200*±*43.390 | *78.830±32.090*** | 159.000*±*26.380 | *76.480±40.220*** |
|  |  | SMI | 2.153*±*0.391 | 2.438*±*0.273 | 2.067*±*0.293 | 2.386*±*0.204 |
|  |  | Tb.Th (mm) | 0.057±0.011 | 0.056±0.009 | 0.058*±*0.007 | 0.057*±*0.007 |
|  |  | Tb.Sp (mm) | 0.216±0.026 | 0.292±0.108 | 0.214±0.007 | *0.330±0.117** |
|  |  | Tb.N | 4.552±0.422 | *3.454±0.633*** | 4.558±0.117 | *3.383±0.995*** |
|  |  | Apparent Density (mg HA/ccm) | 212.800±43.910 | *152.100±30.700*** | 212.500±33.060 | *152.300±32.780*** |
|  |  | Material Density (mg HA/ccm) | 767.600±34.670 | 752.700±29.670 | 765.200±13.030 | 758.800±14.930 |
|  | **Cortical Bone** | BS/BV (mm^-1^) | 21.91±2.731 | *25.08±1.854** | 22.19±1.963 | *24.85±2.519** |
|  |  | BS/TV (mm^-1^) | 8.102±0.6441 | *6.753±0.6802*** | 8.305±0.5110 | *6.798±1.095*** |
|  |  | Porosity (%) | 0.3760±0.05907 | *0.2706±0.03383***** | 0.3776±0.05085 | *0.2750±0.04528***** |
|  |  | Ct.Th (mm) | 0.6241±0.05895 | *0.7295±0.03370***** | 0.6224±0.05085 | *0.7252±0.04516***** |
|  |  | Ct.Sp (mm) | 0.1070±0.01405 | 0.1003±0.01066 | 0.1036±0.009290 | 0.09992±0.007440 |
|  |  | Apparent Density (mg HA/ccm) | 0.2023±0.02073 | *0.2824±0.06401** | 0.2000±0.007714 | *0.2998±0.1018** |
|  |  | Material Density (mg HA/ccm) | 360.800±52.110 | *272.500±32.730***** | 356.700±41.550 | *272.900±42.400**** |
|  |  | BV Density | 835.300±32.010 | 830.400±22.770 | 827.200±15.690 | 824.000±13.570 |

Italics represent measurements demonstrating significant difference from the corresponding sham-vehicle treated group (surrounded by dark border).

(SD: standard deviation; BS: bone surface; BV: bone volume; TV: total volume; CD: connectivity density; SMI: structure model index; Tb: trabecular bone; Ct; cortical bone; Th: thickness; Sp: separation: N: number; mm: millimetre; mg HA/ccm: milligrams of Hydroxyapatite per cubic centimetre; *: p<0.05; **: p<0.01; ***: p<0.001; ****: p<0.0001)
